# Effective connectivity during autobiographical memory search

**DOI:** 10.1101/808501

**Authors:** Norberto Eiji Nawa, Hiroshi Ando

**Author notes:** **Name and full address of corresponding author** Name: Norberto Eiji Nawa, Full address: Center for Information and Neural Networks (CiNet), National Institute of Information and Communications Technology (NICT), Room 2A2, 1-4 Yamadaoka, Suita, Osaka, 565-0871, Japan, Email address, Facsimile.

## Abstract

Autobiographical memory (AM) retrieval is known to recruit a widely distributed network of brain regions but much less is known regarding how these regions interact during the various phases that presumably take place during episodic memory retrieval in general and AM retrieval in particular. Here, we used dynamic causal modeling (DCM) to examine effective connectivity during cued AM search in a sub-network consisting of six major regions within this large network. Functional MRI data was acquired while participants were visually presented verbal cues describing common life events and requested to search for a personal memory that could be associated with each cue. We examined directed couplings between the ventromedial (vmPFC), dorsomedial (dmPFC) and dorsolateral prefrontal cortices (dlPFC), hippocampus, angular gyrus and a region in the posterior midline cortex (RSC/PCC/Prec), all located in the left hemisphere. Results indicated that during AM search, the vmPFC, dlPFC and RSC/PCC/Prec acted as primary drivers of activity in the rest of the network. Moreover, when AM search was completed successfully (Hits), an up-modulation of the effective connectivity of the hippocampus in the vmPFC and angular gyrus was observed. In the same way, there was an increase in the influence of the RSC/PCC/Prec in the activity of the dlPFC and dmPFC. Furthermore, during Hits the angular gyrus showed to have an inhibitory effect in all other nodes of the network. These results are consistent with the notion that midline cortical regions are crucial in supporting the retrieval of AMs, and highlight the interplay between the vmPFC and the RSC/PCC/Prec and the dlPFC during AM search.

## 1. Introduction

Autobiographical memories (AM) retrieval, i.e., when memories of personally experienced events are brought to recollection, is known to engage a large ensemble of brain regions (Cabeza and St Jacques, 2007; Svoboda et al., 2006), most notably the ventral and dorsal aspects of the medial prefrontal cortex, the lateral prefrontal cortex, the posterior medial cortex, likely encompassing portions of the posterior cingulate cortex (PCC), the precuneus and the retrosplenial cortex (RSC), the medial temporal lobes, and the lateral parietal cortex. Even though memories may involuntarily come to mind without conscious effort (Rasmussen and Berntsen, 2011), the large majority of studies so far has typically conceptualized AM retrieval as consisting of a search phase, also known as construction phase (Conway and Pleydell-Pearce, 2000), when a specific memory is searched for guided by an internally or externally generated cue, followed by an elaboration phase, when details associated with the encoding episode are further retrieved and integrated into a vivid construct (Tulving, 1985; Wheeler et al., 1997). The co-activation of this set of regions, or nodes, is now well established, and thought to form a network underlying a variety of different cognitive capacities (Buckner and Carroll, 2007; Spreng et al., 2009; Svoboda et al., 2006). However, studies that have specifically examined the intercouplings within the various regions in that network during AM retrieval are yet to paint a comprehensible picture of the dynamics that takes place during the search and elaboration of autobiographical memories. In (St Jacques et al., 2011), participants were requested to search for AMs associated with auditorily presented emotionally arousing words (both positive and negative), and upon successful recovery of a memory, asked to further elaborate on the retrieved event. Independent component analysis was employed to identify brain-wide spatiotemporal networks (Calhoun et al., 2001) that have been previously linked to different executive functions and top-down cognitive control capacities, such as initiating and adapting task control (frontoparietal network) or task-set maintenance (cinguloopercular network) (Dosenbach et al., 2008; 2007; 2006). Results indicated that networks largely resembling the previously identified frontoparietal and cingulooperculum networks were more associated with the search phase of AM retrieval, whereas a medial prefrontal cortex network and a medial temporal lobe network were equally associated with both the search and elaboration phases. The same study investigated effective connectivity during search and elaboration using dynamic causal modeling (DCM) (Friston et al., 2003), albeit by means of a rather unusual approach (Stevens et al., 2007), where instead of examining couplings between spatially circumscribed regions of interest (ROIs), as typically done in most DCM studies, they examined couplings between the large networks found to be associated with the search and/or elaboration phases. DCM allows one to assess how the nodes within a network are connected to each other (or in the aforementioned study, how networks comprising several regions are connected to other networks), the directions and magnitudes of the connections, i.e., which nodes effect the activity in other nodes and by how much, as well as the valences of such connections, i.e., whether excitatory or inhibitory. Results indicated that a medial prefrontal cortex network – consisting of dorsomedial prefrontal cortex (dmPFC), PCC, and ventral parietal cortex – drove the activation in the other networks during AM retrieval, in both the search and elaboration phases. Interestingly, results also pointed out to the existence of a medial temporal lobe network, encompassing regions that are typically associated with memory retrieval processes such as ventromedial prefrontal cortex (vmPFC), hippocampus, and parahippocampus, which influenced the medial PFC network during memory search but only in the trials where the retrieved AM was more *accessible* (i.e., when participants were able to quickly find a memory associated with the cue). These results highlight first and foremost the involvement of widely distributed brain networks during the performance of AM retrieval. Nevertheless, it is not clear why the medial PFC network, and not the network containing regions more closely associated with memory retrieval processes (medial temporal lobe network) or the networks strongly associated with executive control functions (frontoparietal and cingulooperculum networks), was found to primarily drive the activity in the other networks during both memory search and elaboration. In addition, how regions within and between such large networks interact with one another during AM retrieval processes still remains to be clarified.

McCormick et al. (2015) applied multivariate statistical analysis to examine whether there are changes in functional and effective connectivity between the hippocampus and the rest of the cortex when transitioning from AM search to AM elaboration. Because there appears to be functional and connectivity-wise distinctions between the anterior and posterior hippocampus (Dalton et al., 2019; Zeidman and Maguire, 2016), effective connectivity was examined based on independent voxels from both hippocampal subregions (bilaterally), plus regions that were found to be functionally connected with a seed located in the left anterior hippocampus, namely, the left dmPFC, the left ventrolateral PFC, the left medial PFC, the middle occipital cortex (bilaterally), the left lingual gyrus and the right fusiform gyrus. Structural equation modeling analysis based on timeseries extracted from these eleven voxels primarily revealed distinct effective connectivity of anterior and posterior hippocampus with the selected cortical regions, during both AM search and AM elaboration; the left anterior hippocampus was found to exert greater positive influence in the dorsomedial PFC and right anterior hippocampus during AM search than during AM elaboration; on the other hand, the posterior hippocampus (bilaterally) was found to exert greater influence in the middle occipital and fusiform gyrus during AM elaboration than during AM search. These results suggest that AM search may be characterized by a greater (anterior) hippocampus to (dorsomedial) PFC connectivity, whereas during AM elaboration, the effect of the (posterior) hippocampus majorly shifts to regions that are typically associated with visual processing. Though there is still no consensus regarding the degree of involvement of the hippocampus in the neurophysiological mechanisms underlying episodic memory retrieval in general (Nadel et al., 2007; Squire and Bayley, 2007), more recent views have argued for a shift in the focus of the discussion from the structures located in the medial temporal lobe, in particular the hippocampus, to one that emphasizes the interactions between such structures and the prefrontal cortex (Eichenbaum, 2017; McCormick et al., 2018; Rubin et al., 2017). Even though the results in (McCormick et al., 2015) pointed out to a heightened hippocampal-dmPFC interaction during AM search, such effects were not observed with the left medial PFC, the node most likely to be located in the vmPFC at large, an area that has been strongly linked with AM retrieval processes (Bonnici et al., 2012; McCormick et al., 2018; Nieuwenhuis and Takashima, 2011). Also, it is worth noting that the regions assessed in the effective connectivity analysis were selected based on the degree of functional connectivity with a left anterior hippocampus seed. That biased procedure possibly overlooked regions that were not temporally in lockstep with the anterior hippocampus but still play relevant roles during the processes underlying the retrieval of AMs, in concert or in parallel with the hippocampus.

To further extend this growing body of research, using functional MRI (fMRI), we applied DCM to examine effective connectivity in a network composed by 6 major regions that have been consistently shown to co-activate during AM retrieval processes, namely, the vmPFC, dmPFC and dorsolateral (dlPFC) prefrontal cortices, hippocampus, angular gyrus and the posterior midline cortex. Because the goal was to characterize the couplings that take place specifically during cued AM search, our experimental task deliberately did not include an AM elaboration phase. AM search is thought to rely on an effortful process of iterative search through an autobiographical knowledge base (generative retrieval) that starts with the recovery of highly abstract knowledge about the self, and is followed by a process involving iterations through search cycles that gradually refines the recovered knowledge, finally culminating in the retrieval of a specific event that fulfills the original search criteria (Conway, 2005; Conway and Pleydell-Pearce, 2000; Haque and Conway, 2001). Generative retrieval has been shown to preferentially recruit lateral prefrontal and temporal regions, possibly reflecting strategic and executive control processes associated with memory search operations (Addis et al., 2012). A crucial question regarding the dynamic interaction between these regions during retrieval is, naturally, the direction of such influences. For instance, with regard to the vmPFC and hippocampus, the evidence from studies focusing on the construction of imaginary events is so far mixed, with results showing both enhanced effective connectivity from the hippocampus to the vmPFC (Campbell et al., 2017), as well as in the reverse direction, from the vmPFC to the hippocampus (Barry et al., 2019a). Here, we hypothesized that during AM search, prefrontal regions would predominantly influence the activity in the rest of the network, including the hippocampus, primarily due to the involvement of lateral and medial prefrontal regions in executive control processes and episodic memory-specific processes, Furthermore, we hypothesized that activity in prefrontal regions would be inhibited in trials where a memory was successfully found, though we did not have specific hypotheses about which region (or regions) would be in the acting end of such an effect.

## 2. Material & Methods

### 2.1 Participants and study design

Forty-three right-handed volunteers, all fluent Japanese speakers, were initially recruited to this study (22 females, mean age 22.6 years, range 20-27) via a part-time employment agency. We limited the age of the participants to the 20-30 years old range, in order to promote some uniformity in terms of the age of the memories recalled during the experiment across participants. All participants gave informed written consent prior to participation in the experiments, in accordance with the principles stated in the Declaration of Helsinki. The study was approved by the local research ethics committee. All participants had normal or corrected-to-normal vision and declared that they were not receiving treatment for psychiatric disorders at the time of the study and had no history of neurological diseases (one participant declared having received medication prescribed by a psychiatrist in the past). Before entering the scanner, participants completed the Beck Depression Inventory (BDI-II) (Beck et al., 1996; Kojima et al., 2002), the Edinburgh Handedness Inventory (Oldfield, 1971) and the Positive and Negative Affective Scale (PANAS) (Watson et al., 1988). The PANAS was collected again after participants exited the scanner, along with the Vividness of Visual Imagery Questionnaire (VVIQ) (Marks, 1973). The VVIQ was scored using a reversed scale to allow for comparisons with a previous report (Zeman et al., 2015).

All participants underwent scanning and were monetarily compensated for their time. One participant was unable to complete the task scanning sessions due to technical problems, 1 participant displayed an anatomical abnormality in the right temporal pole, and 4 participants had a BDI-II score greater than 12 (a screening level adopted in other studies, e.g., (Leal et al., 2014; Nawa and Ando, 2019)); their data were excluded from the analyses upfront, resulting in an initial cohort of *N=*37 participants (19 females, mean age 22.4 years, range 20-27, mean BDI-II score 4.2, mean handedness laterality coefficient 88.2%).

### 2.2 FMRI experimental paradigm

Participants performed an AM search task (Fig. 1) inside the scanner. Each trial started with a fixation period of 8-10 seconds (mean of 9 s), which was immediately followed by the display of a verbal cue (“Trip with a friend”) on the screen (20 s). Participants were instructed to look for an AM that could be somehow associated with the cue; they were additionally told that the retrieved memory did not have to be a perfect match. An autobiographical memory was defined as a memory associated with a specific event that they themselves had experienced in the past and should necessarily be characterized by a specific time and place of occurrence. If they could find such a memory, participants were instructed to press the button corresponding to the side (left or right) where the choice “Yes” was displayed on the screen in that trial; if they were unable to find an appropriate memory within the allocated time, participants were told to press the opposite button (“No”). The sides in which the choices appeared on the screen were randomized across trials and participants; each choice appeared the same number of times on either side. After the button press, participants were instructed to relax and wait for the next trial; if they had pressed “Yes”, they were additionally asked not to purposefully engage in thoughts associated with the retrieved memory, i.e., elaborate the memory. All text was presented in white against a black background (opposite to the schematic in Fig. 1). During the last 3 seconds of each trial the color of the verbal cue was changed to purple to signal participants that the end of the trial was approaching; they were requested to necessarily make a decision before the end of the trial. The order of presentation of the verbal cues was randomized across participants. No constraints were imposed with regard to the age of the retrieved memories.

**Figure 1:**
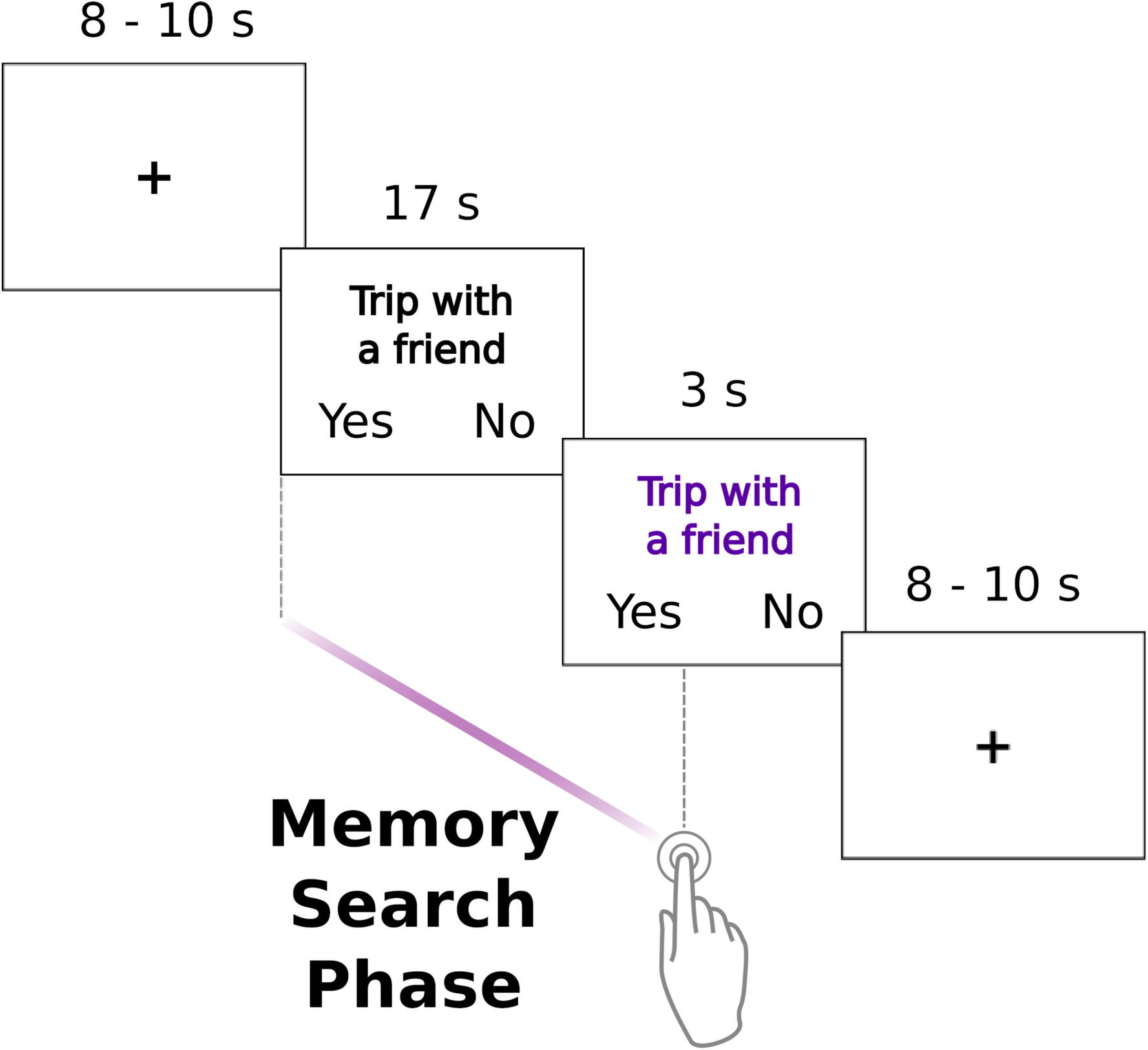
One trial of the AM search task. Participants were instructed to press the button corresponding to “Yes” as soon as they were able to find a memory that could be associated with the verbal cue (“Trip with a friend”), otherwise, they should press the opposite button. The color of the cue changed to purple during the last 3 seconds of each trial.

Twelve verbal cues were presented in each one of the 6 scanning sessions. Each session lasted 362 seconds. Participants completed all scanning sessions in the same day, and they were encouraged to take short breaks between sessions. The AM search task was implemented using the software Presentation v.18.2, (http://www.neurobs.com). Images were projected onto a screen located outside the bore, and participants viewed the screen through a mirror mounted on the head-coil. Participants were told beforehand that upon leaving the scanner, they would be asked to review again each one of the cues presented during the fMRI experiment. For the verbal cues that could be associated with a personal memory during scanning, participants were asked to classify the memory as a *positive*, *negative* or *neutral* (neither positive nor negative). Furthermore, they were asked to evaluate the (1) magnitude of the positive affect elicited when recalling that memory, (2) the overall vividness of the imagery evoked when recalling that memory, (3) the intensity of the emotional response experienced during the encoding event, (4) the personal significance of that event, (5) the effort that was necessary to retrieve the associated memory, and (6) their age at the time of occurrence. Ratings for questions 1) to 5) were given in a 4-point scale with 1: low to 4: high. Participants were also requested to write short sentences describing each one of the events; those sentences served as stimuli material in a subsequent study that will be reported elsewhere.

### 2.3 Verbal cues

Verbal cues were 72 short sentences describing typical life events, and were selected from a list of 110 cues employed in a previous study (Nawa and Ando, 2019) as follows: Cues that were commonly associated with an autobiographical memory but often evocative of extreme negative emotions were removed from the list; the remaining cues were then ranked in terms of “popularity”, i.e., the likelihood of a cue being associated with an AM based on responses given by a different sample of 44 individuals, and the top 72 cues were selected to be used in this study. Of the selected cues, the most and least “popular” cues were associated with an autobiographical memory by 97.7% and 45.5% of the individuals in the testing sample, respectively. Personally-relevant cues or cues that are familiar to the participants may provide a direct point of entry to a specific AM (Conway and Pleydell-Pearce, 2000) (direct retrieval), thus requiring considerably less effort. To prevent any beforehand preparation, participants in this study were first exposed to the cues only during scanning, and a second time when performing the post-scan ratings.

### 2.4 Imaging data acquisition

A 3T Siemens Magnetom Trio whole-body MR scanner equipped with a standard 32-channel head-coil was used to acquire the imaging data. Participants entered the scanner after being screened for MRI contradictions, and briefed on MR safety and general procedures. They wore earplugs to attenuate scanner noise, and hand towels were used to fill in the space between the head and the coil in order to minimize head movement and minimize discomfort. First, a standard double-echo gradient echo field map sequence images were collected for distortion correction of the functional images [echo time (TE1) = 4.92 ms, TE2 = 7.38 ms, voxel size = 2. 0 mm^3^, repetition time (TR) = 739 ms, flip angle = 90°]. Next, resting-state data (no experimental task) were collected over the course of a single session (203 functional images using a T2*-weighted multiband (Moeller et al., 2008) echo planar imaging sequence (EPI), TR = 2000 ms; TE = 30 ms; flip angle = 75°; field of view (FOV) = 200 mm; voxel size = 2.0 mm^3^ isotropic; 75 axial slices; acceleration factor 3). Slices were posteriorly tilted approximately 20 degrees off the AC-PC line to minimize signal dropout near the ventral medial prefrontal cortex and the orbital sinuses. Participants were instructed to close their eyes but to keep awake and avoid continuously thinking about something specific. Resting-state data were acquired prior to the task sessions to prevent any potential contamination from activity associated with the performance of the AM search task. Due to technical problems during scanning, we were unable to collect resting-state data from 3 participants. Following the resting-state session, a whole-brain T1 MPRAGE anatomical image was acquired for coregistration and normalization purposes [1.0 mm^3^ isotropic, flip angle = 9°, TR = 1900 ms, time for inversion (TI) = 900 ms, TE = 2.48 ms]. The anatomical scan lasted approximately 4 minutes; during that acquisition, participants practiced the AM search task using specially prepared extra cues. Following the anatomical scan, 182 whole-brain EPI functional images were acquired in each one of the 6 sessions on which participants executed the AM search task, using the same acquisition parameters employed in the resting-state session. Participants held a response box (4-button, diamond layout, by Current Designs, http://www.curdes.com) in their right hands to record behavioral responses (button presses were done using the right thumb), and a squeeze ball for emergency purposes in their left hands. Task sessions where excessive head movement was detected (peak translation in any one direction > 2 mm) were excluded from the analyses (2 participants, one session each).

### 2.5 Imaging data processing

Imaging data from the task sessions were processed and analyzed using Statistical Parameter Mapping (SPM12, v7487, Wellcome Trust Centre for Neuroimaging, London, UK, RRID:SCR_00703). The first 3 images of each task session as well as the resting-state session were discarded to allow for magnetic field stabilization. The functional images from the task and resting-state sessions were first corrected for geometric distortions using the field maps. They were then spatially realigned within and across sessions using a rigid body transformation to correct for head movement (the first image of each session, and the first image of the first session used as references) and unwarped in order to correct for gradient-field inhomogeneities caused by motion. From this step onward, data from the resting-state session was analyzed using the toolbox CONN (version 18.b, https://www.nitrc.org/projects/conn, RRID: SCR_009550) (Whitfield-Gabrieli and Nieto-Castanon, 2012), where the rest of the preprocessing and analysis were performed (see Supporting Information). For the analysis of the data collected during the task sessions, the T1 anatomical image of each participant was coregistered to the mean functional image generated after realignment/unwarping. Task-based functional images were normalized to the MNI template space by applying parameters derived from the normalization of the participant’s T1 anatomical image to the MNI/ICBM template (East Asian brains). The normalized images were rewritten at 2 mm isometric voxels, and spatially smoothed with a 6 mm full-width half-maximum (FWHM) Gaussian kernel.

### 2.6 Behavioral Data

We examined whether there were differences in terms of reaction time (RT) between trials where a memory associated with the verbal cue was successfully found (Hits) from trials where participants were unable to find a memory within the allotted time (Misses). For each participant, the mean time elapsed between the verbal cue onset and a button press was computed for both trial types, and the data were entered in a two-sided paired samples Wilcoxon signed-rank test. Trials in which button presses were not recorded were excluded from the analysis. We also assessed the characteristics of the memories associated with the cues by examining the data from the post-scan questionnaires.

### 2.7 Examining effective connectivity in the AM retrieval network during memory search using DCM

#### 2.7.1 First-level analysis

A mass-univariate analysis was performed to identify brain regions recruited during AM search. First-level general linear models (GLMs) were computed using the normalized and spatially smoothed images. Trials were classified based on the button presses given by the participants; brain activity recorded during AM search was modeled as a boxcar function starting at the onset of the verbal cue and ending with the button press, using two regressors (Hits, Misses). Trials in which button presses were not recorded were modeled using a separate regressor (No response). Fixation screens and button presses were not entered in the GLMs. To generate the predicted blood-oxygenation level dependent (BOLD) responses, the boxcar functions were convolved with the canonical hemodynamic response function implemented in SPM. Six head movement parameters derived from the realignment step were incorporated as regressors of no interest. An autoregressive AR(1) model was used to correct for timeseries correlations during model parameter estimation, and a highpass filter (cutoff 128 s) was applied to remove slow signal drifts. First-level contrasts were computed based on the resulting voxelwise parameter estimates; we computed the contrast [Hits], to verify whether the brain regions recruited during successful AM search were consistent with previous reports, and the contrast [Hits + Misses] to determine the group-level peak voxels that would serve to guide the extraction of the timeseries data used in the DCM analysis.

#### 2.7.2 Group-level analysis

The first-level contrasts were entered into a group-level analysis with participant as a random factor, and whole-brain voxelwise one-sample t-tests were performed. A family-wise-error correction for multiple comparisons (FWE) for the whole-brain implemented in SPM, at a height threshold of *p <* 0.05, was adopted to determine the brain regions that displayed enhanced activity relative to the implicit baseline.

#### 2.7.3 Extracting individual timeseries data

We used the coordinates of the group-level peak voxels of each one of the 6 ROIs to first identify individual-level peak voxels. For each one of the participants, we examined the results of the first-level contrast [Hits + Misses], using a threshold of p < 0.005 (uncorrected), and looked for local maxima within the voxels contained in 5 mm-spheres centered at each one of the group-level peak voxels. Participants should have at least one supra-threshold voxel within each one of the 6 spherical regions to be included in the DCM analysis; participants that did not satisfy this criterion were left out from the analysis. Target regions for data extraction were defined as 5 mm-spheres centered at the newly found individual-level peak voxels. The SPM Volume of Interest utility was employed to extract timeseries data from the target regions. A liberal threshold of p < 0.05 (uncorrected) was employed to determine supra-threshold voxels within each target region, in line with other studies, e.g., (Fastenrath et al., 2014). Timeseries were extracted from normalized, but not spatially smoothed, data as the first eigenvariate across supra-threshold voxels with the target region, and were adjusted for ‘effects of interest’, i.e., they were mean-corrected and rectified based on the movement parameters obtained after spatial realignment.

#### 2.7.4 DCM analysis

Dynamic Causal Modeling (DCM) (Friston et al., 2003) is a method for estimating effective connectivity across brain regions, or nodes, i.e., how the neural activity in one brain region effects activity in another brain region (Friston, 2009). In a typical situation, there will be several candidate-models reflecting different hypotheses about the characteristics of the network underlying a given cognitive function or perceptual process. DCM can determine the model that most parsimoniously explains the observed data among the assessed models, if there is one, providing a principled way to compare different hypotheses regarding the directions, strengths and valences (whether excitatory or inhibitory) of the network connections, and thus, enabling inferences about the organization of the underlying functional brain network. Under the same framework, DCM also allows one to examine how external modulatory effects influence the strength of connections, i.e., whether and how experimentally controlled manipulations alter the effective connectivity exerted by one node onto another. A recent addition to the DCM array of tools was the introduction of the Parametric Empirical Bayes (PEB) framework (Friston et al., 2016; Zeidman et al., 2019a; 2019b), which allows the efficient identification of commonalities across different models generated from the same participant, or most importantly, across participants. Under the PEB framework, only one comprehensive model has to be computed for each participant; the model parameters are then taken to a group-level analysis, where they can be examined across participants, much along the spirit of first-level and group-level mass-univariate GLM analysis. Based on the first-level estimation results of individual fully-connected DCMs, a search over the nested parameter space is performed at the group-level (also known as post-hoc search or Bayesian Model Reduction), enabling one to determine the model parameters that do not contribute to model evidence, and hence, should not be included in the “minimal” model. Here, for each participant we estimated a bilinear, deterministic, one-state, fully-connected DCM with mean-centered inputs, where all nodes were connected to every other node, all nodes received the external driving input (which consisted of the onsets, with the respective durations, of all valid AM search trials, in effect, the union of the Hit trials and the Miss trials), and all connections, including the self-connections, were subject to the external modulatory input (which consisted of the onsets, with the respective durations, of the Hit trials). We only report the parameters that had a posterior probability equal or above 95%. Using such an approach, one can determine the most likely node, or group of nodes, whose activity is primarily driven by the external driving input (operationalized as the onsets of all Hit and Miss trials), and the underlying organization of the 6-node network during AM search, i.e., the directions of the connections, their strengths and valences, and the connections that are up- or down-modulated in the trials where a personal memory was successfully found (operationalized as the Hit trials). The strength of the directed between-region endogenous connections in the network are rates of change, i.e., they indicate how much the activity in the node receiving the connection is effected by the activity in the node sending the connection. The strength of such connections is represented in units of Hz, and they can be excitatory, meaning that the sending node increases activity in the receiving node (positive effect), or inhibitory, meaning that the sending node decreases activity in the receiving node (negative effect). Under the DCM framework, the self-connections (*A_self_*) must be negative by definition so they are treated as unit-less log-scaling parameters, and converted to rates of change (*a*) when necessary using the following equation, *a* = -exp(*A*_self_)*0.5.

Here, all analyses were performed using functions provided with SPM 12 (release 7487, DCM12). Nodes employed in the DCM analysis were restricted to the left hemisphere because, though evidence of a clear lateralization regarding AM retrieval processes is still mixed especially with regard to the hippocampus (Piefke et al., 2003; Ryan et al., 2001; Viard et al., 2007), previous studies have reported the predominant involvement of left-lateralized regions (Addis et al., 2004; Cabeza and St Jacques, 2007; Conway et al., 1999; Gardini et al., 2006; Gilboa, 2004; Maguire, 2001; Maguire and Frith, 2003; Maguire and Mummery, 1999; Piolino et al., 2009; Svoboda et al., 2006).

## 3. Results

### 3.1 Behavioral data

Though a certain variability in the distribution between Hits and Misses across participants was naturally expected to be observed, to prevent the inclusion of extremely unbalanced participants (which could potentially affect the reliability of the parameter estimates), we established an arbitrary cutoff criterion of a minimum of 20% of the valid trials having to be either Hits or Misses. Based on that criterion, upon inspection of the behavioral responses given inside the scanner, 13 participants had to be dropped from the sample resulting in a subset of 24 participants (10 females, mean age 22.6 years, range 20-27, mean BDI-II score 4.6). The average occurrence and range of Hits among the included and excluded participants was 65.6% [33.3% - 79.2%], and 88.0% [80.6% - 95.8%], respectively, indicating that the excluded participants displayed higher success rates when searching for a personal memory that could be associated with the verbal cues.

We examined RT differences between Hit trials and Miss trials based on the responses given by the 24 participants. Individual mean RTs for Hit and Miss trials were entered in a Wilcoxon signed-rank test. Results indicated that participants responded significantly faster in Hit trials (mean = 7.38 s, range [3.09 s – 12.79 s]) than in Miss trials (mean = 8.78 s, range [4.35 s – 17.22 s]), Z = −2.97, p = 0.030. Inspection of the RTs for Miss trials using a histogram (20 bins) revealed the existence of a relatively large concentration of occurrences just around 17 seconds after the trial onset, suggesting that participants often decided that they had no memory associated with the cue right after being signaled that there were only 3 seconds remaining in the trial (Fig. S1, Supporting Information).

Across the same cohort, the (reversed) mean VVIQ collected after scanning was 53.5 (range 36 - 68), which majorly overlaps with the range of values reported by normal participants (Zeman et al., 2015). No differences were detected in the Positive or Negative Affect Scale scores collected before and after scanning (Wilcoxon signed-rank test, p = 0.625, and p = 0.250, respectively). Participants diligently performed the AM search task; the average rate of no-response trials across participants was only 0.46%.

Results from the post-scan questionnaires indicated that across participants, cues were associated with memories that were almost equally likely to be classified as *positive* (44.6%, range [26.5% - 61.1%]) or *neutral* (43.9%, range [21.6% - 65.9%]), likely reflecting the criterion used to pre-select the verbal cues. Despite the fact that we filtered out cues that were likely to be associated with extreme negative memories, 11.5% of the retrieved memories were classified as *negative* (range [3.7% - 32.4%]). Responses to the other questions regarding the retrieved memories are summarized in Table 1.

**Table 1:** Mean responses (with respective range of values across participants) given to the questions about the retrieved memories collected in the post-scan questionnaire (24 participants). Responses were given using a scale from 1: low to 4: high.

|  | Mean | Range |
| --- | --- | --- |
| Positive affect elicited when recalling the memories | 2.3 | 1.5 – 3.0 |
| Vividness of the memory imagery | 2.8 | 1.9 - 3.7 |
| Emotional intensity experienced during the encoding event | 2.6 | 1.9 – 3.4 |
| Personal significance of the event | 2.2 | 1.5 – 2.3 |
| Effort necessary to retrieve the memory | 1.8 | 1.1 – 2.9 |
| Age of the memories | 2.8 | 0.1 – 7.5 |

### 3.2 Mass-univariate general linear model analysis

The group-level results for the contrast [Hits] are shown in Figure 2, highlighting the brain regions that displayed enhanced activation during the Hit trials, compared to the implicit baseline. Activity in areas typically associated with memory retrieval processes was observed spanning over a wide network. Salient clusters were identified in the lateral prefrontal cortex, as well as in the lateral parietal cortex, predominantly in the left hemisphere. In cortical midline regions, we observed clusters of enhanced activity in both prefrontal and posterior regions, including the ventromedial and dorsomedial aspects of the prefrontal cortex, and the posteriomedial cortex. There were clusters of activity in the medial temporal lobes (MTL), including hippocampus and parahippocampal cortices, both bilaterally. We verified the existence of supra-threshold voxels, as well as clusters of activity in all left-lateralized 6 ROIs at a p < 0.05 (FWE) using this contrast (data not shown).

**Figure 2:**
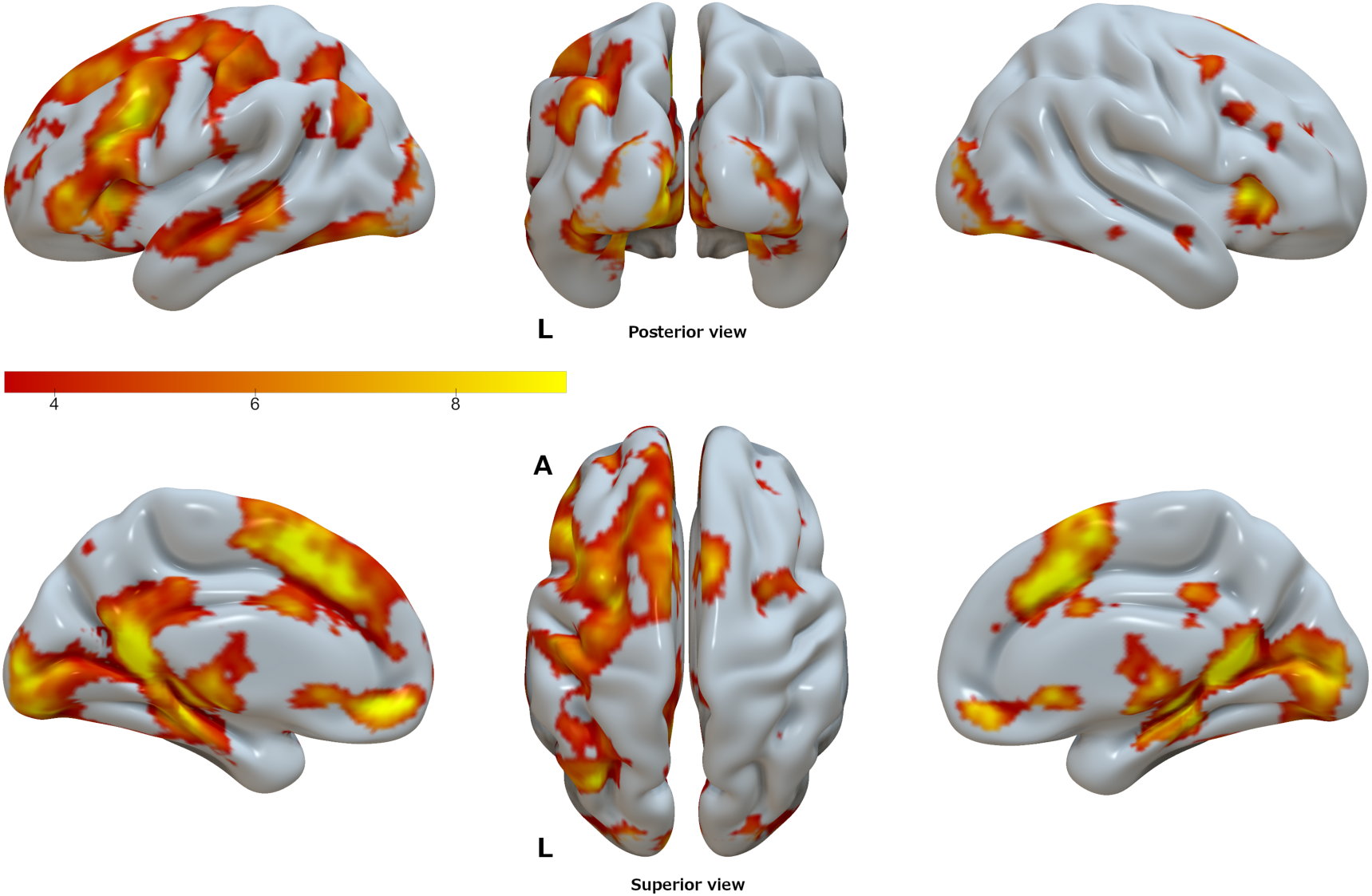
Results of the mass-univariate GLM analysis showing the areas in the brain that displayed enhanced activity during the Hit trials, compared to the implicit baseline; T-values are overlaid on a semi-inflated smoothed version of the ICBM152 brain using Surf Ice. Results are shown at p < 0.001 uncorrected, for illustration purposes. Left panels show the lateral (top) and medial (bottom) views of the left hemisphere. Right panels show the corresponding data for the right hemisphere. Middle panels show the posterior (top) and superior (bottom) views of both hemispheres.

After confirming that the AM search task recruited regions typically involved in AM retrieval processes, we assessed the group-level results generated by the contrast [Hits + Misses] at a p < 0.05 (FWE), and again specifically looked for clusters of activity in the vicinity of the 6 ROIs. The group-level peak voxels are shown in the Table 2 and rendered in Figure 3. Because there is still much debate about the functional organization of the posterior medial cortex, and to avoid any premature (mis)labelling of the region that was encountered, we deliberately opted to use a comprehensive label to cover this region by combining three labels commonly assigned to this area in episodic memory studies, namely, retrosplenial cortex (RSC), posterior cingulate cortex (PCC) and Precuneus (Prec).

**Figure 3:**
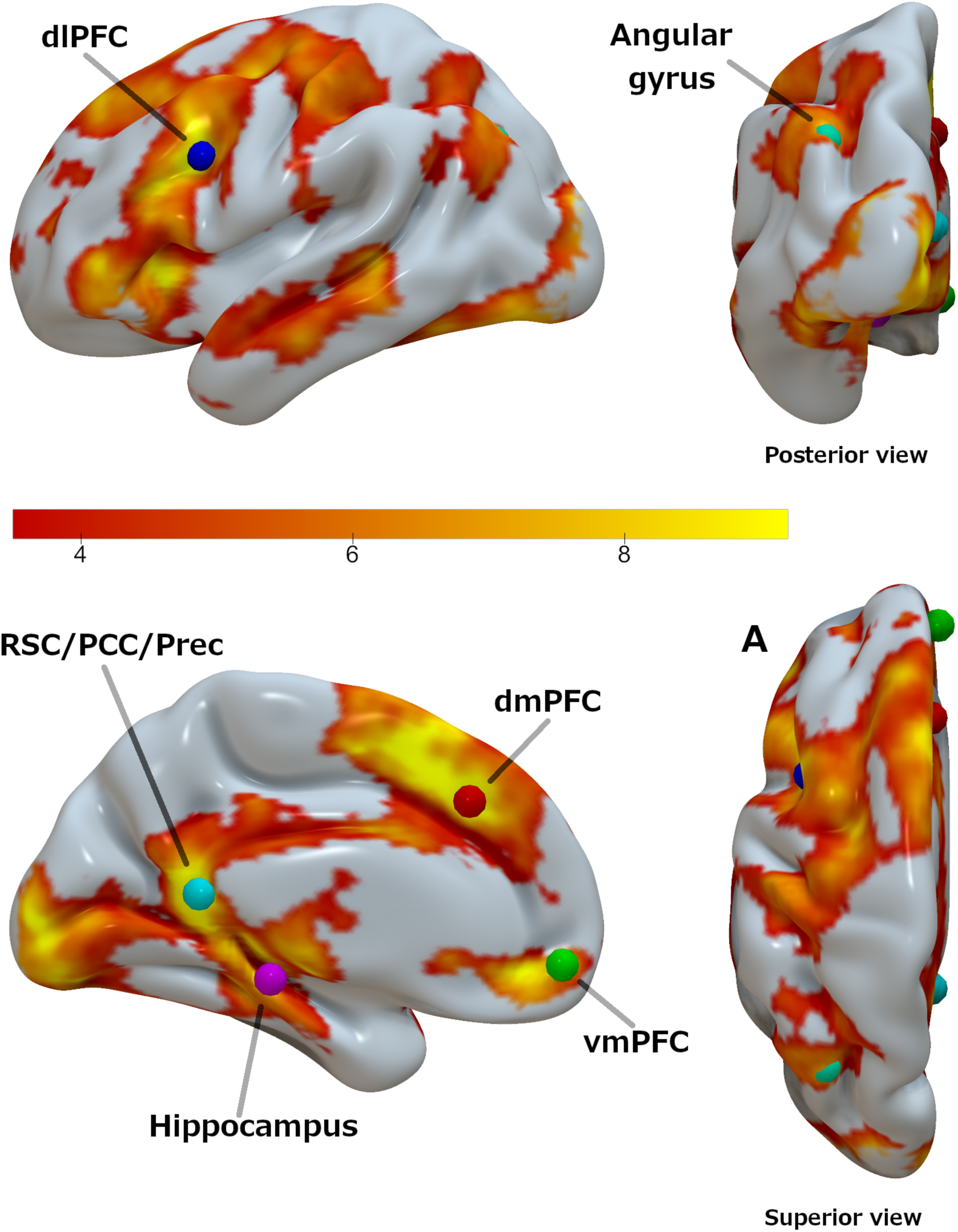
Group-level results of the mass-univariate GLM analysis for the contrast [Hits + Misses], compared to the implicit baseline; T-values are overlaid on a semi-inflated smoothed version of the ICBM152 brain using Surf Ice. Results are shown at p < 0.001 uncorrected, for illustration purposes. Colored dots are 5mm-spheres centered at the group-level peak voxels of the 6 ROIs used to guide the extraction of the timeseries used in the DCM analysis.

**Table 2:** MNI coordinates and statistical results for the group-level peak voxels in the 6 ROIs, determined using the contrast [Hits + Misses]; p-values corrected for multiple comparisons (family-wise error correction on a whole-brain level). RSC/PCC/Prec: posterior medial cortex, dmPFC: dorsomedial prefrontal cortex, dlPFC: dorsolateral prefrontal cortex, vmPFC: ventromedial prefrontal cortex, L HPC: left hippocampus.

|  | x | y | z | T | Z | p |
| --- | --- | --- | --- | --- | --- | --- |
| RSC/PCC/Prec | -6 | -48 | 12 | 12.69 | 6.85 | 0.000 |
| dmPFC | -6 | 28 | 38 | 12.10 | 6.72 | 0.000 |
| dIPFC | -40 | 12 | 32 | 12.06 | 6.71 | 0.000 |
| vmPFC | -4 | 54 | -8 | 9.53 | 6.01 | 0.000 |
| L HPC | -22 | -28 | -12 | 9.24 | 5.92 | 0.000 |
| Angular gyrus | -36 | -70 | 36 | 8.74 | 5.75 | 0.001 |

### 3.3 DCM

When examining the 5 mm vicinity around the group-level peak voxels on the results of the first-level contrast of each participant, we were not able to verify the existence of supra-threshold voxels in all 6 ROIs in the data from 3 participants, resulting in a final group of *N=*21 participants (9 female, mean age 22.2 years, range 20-26, mean BDI-II score 4.7). Individual fully-connected DCM models were computed, and the search over the nested parameter space (PEB) was performed at the group-level. The nested search identifies model parameters of the fully-connected DCM that do not relevantly contribute to the model evidence, by switching each parameter on and off, and examining the resulting differences in model evidence with regard to a posterior probability (Pp) threshold. We first examined the results regarding the location of the external driving input (in DCM jargon, the matrix C), i.e., which nodes had activity directly driven by the onsets of the main experimental manipulation of the AM search task. In the context of DCM, such nodes can be interpreted as the points from where activity is initiated, before it reverberates to the rest of the network. Here, the dlPFC was found to be the sole location of the external driving input, even surviving a stricter threshold of Pp > 99%.

We then examined the endogenous connections between the 6 ROIs, including the self-connections (in DCM jargon, the matrix A). The endogenous connections represent the average effective connectivity strength across all experimental conditions, which in this case correspond to the periods of time when people performed the AM search task after being cued, i.e., regardless of whether the trial resulted in a Hit or Miss. Results showed first and foremost that the 6-node network was almost fully interconnected by excitatory (positive) and inhibitory (negative) connections (Fig. 4), with the only exceptions being the angular gyrus to dlPFC and vmPFC connections (though the same connections were down-modulated during Hit trials), and the dmPFC to hippocampus connectivity being unsupported at all. The dlPFC had positive connections to all other nodes in the network; similarly, the vmPFC had positive connections to all other nodes but the angular gyrus. The RSC/PCC/Prec had positive connections to the vmPFC and the angular gyrus, but otherwise all other connections were negative. The dlPFC and the vmPFC both sent and received positive connections from each other, in effect, constituting a positive loop during AM search. A similar loop was observed between the vmPFC and the RSC/PCC/Prec. Conversely, connections leaving the hippocampus, angular gyrus, and dmPFC were all negative or unsupported, putting them in a counterbalancing role with regard to the excitatory effects brought by the other nodes.

**Figure 4:**
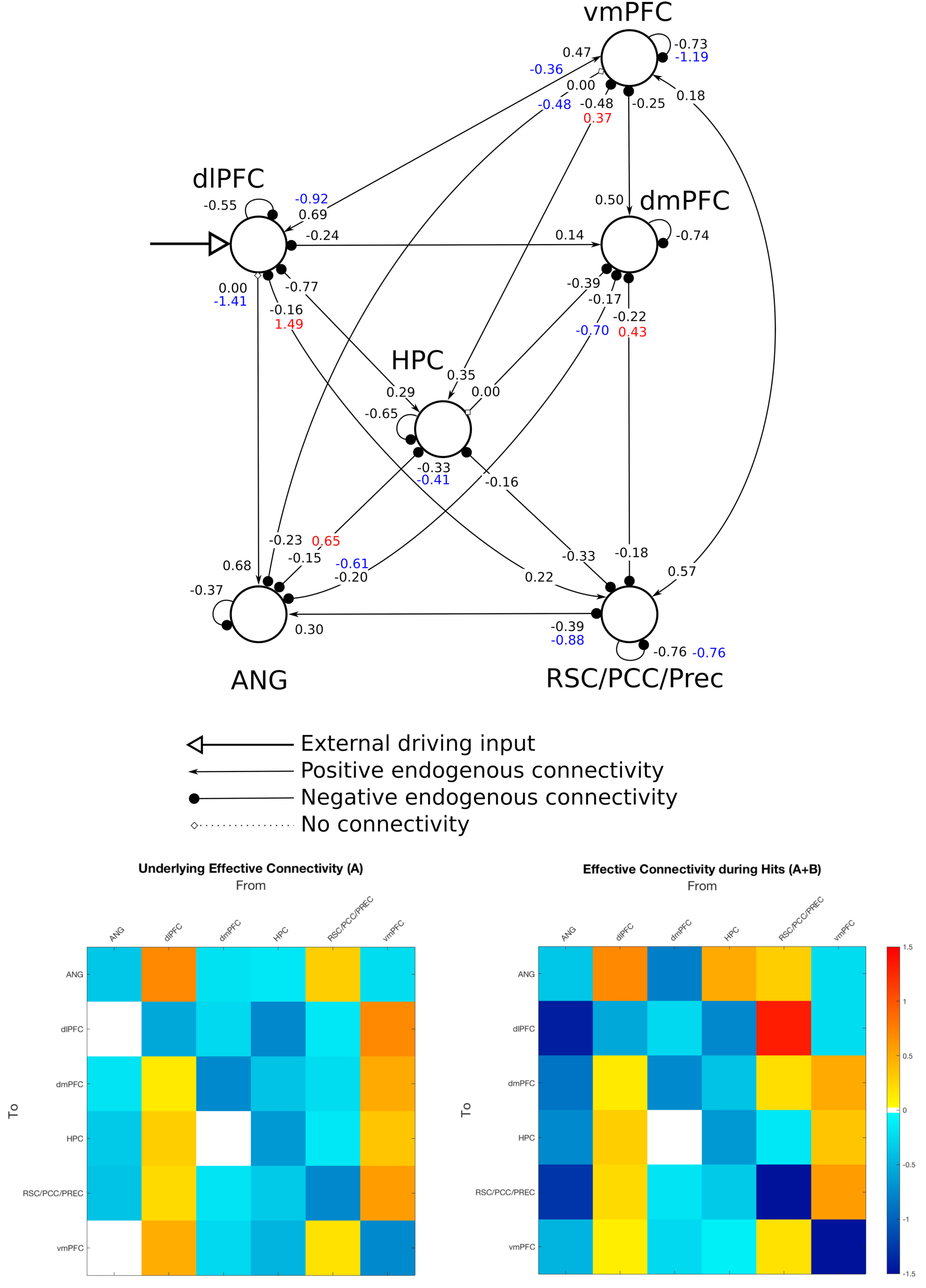
Results of the DCM analysis. (Top panel) The strength of the endogenous connectivity is displayed near the receiving end of the link. All coupling parameters are in units of change rates (Hz), with the exception of the self-connecting parameters, which are log-scaling parameters. For the connections that were subject to modulatory effects associated with the Hit trials, the strength of the modulatory effect is displayed near the respective endogenous connectivity strength in red (excitatory) or blue (inhibitory). (Bottom panel) Matrix representation of the same results showing the magnitude of the endogenous connectivity (left), i.e., the matrix A in DCM jargon, and the total effective connectivity during Hits (right), i.e., the sum of the endogenous connectivity with the modulatory effect (in effect, the sum of matrices A and B). Positive (excitatory) connections are shown in yellow-red, negative (inhibitory) connections are shown in cyan-blue.

Finally, we looked at the modulatory effects associated with the Hit trials in the 6-node network (in DCM jargon, the matrix B). Modulatory effects act on top of the endogenous connections, so during Hit trials, the net effective connectivity amounts to the sum of both values. The connections departing from the RSC/PCC/Prec and arriving at the dlPFC and dmPFC, which were both originally negative on average across trials, were positively modulated during Hit trials. In particular, the RSC/PCC/Prec-to-dlPFC connectivity displayed the greatest magnitude among all connections in the network. Connections from the hippocampus to the vmPFC and also to the angular gyrus were also positively modulated during Hit trials. In contrast, the connections linking the dlPFC to the vmPFC in both directions were negatively modulated. One remarkable finding was that connections from the angular gyrus to all other nodes – including the links to the dlPFC and the vmPFC, which were originally unsupported – were negatively modulated during Hit trials. In a similar manner, the connection from the dmPFC to the angular gyrus was also negatively modulated during Hit trials. Finally, the self-connections in the vmPFC and RSC/PCC/Prec were also negatively modulated during Hits. All results from the DCM analysis were initially assessed at the Pp > 95% level but remained identical at the Pp > 99% level as well.

## Discussion

The present study used DCM to assess effective connectivity during AM retrieval in a network formed by 6 brain regions that are thought to be part of a larger “core” network supporting, among other things, the processes underlying the retrieval of episodic memories (Cabeza and St Jacques, 2007; Schacter et al., 2012; Svoboda et al., 2006). Here we focused the analysis in regions along the frontal (vmPFC, dmPFC) and posterior (RSC/PCC/Prec) midline cortices, which mostly overlap with the default-mode network, regions involved with attention and cognitive control (angular gyrus and dlPFC), and the hippocampus. DCM results (Section 3.3) showed that the RSC/PCC/Prec, dlPFC and vmPFC were the only nodes that positively influenced the activity in the rest of the nodes via endogenous connections, suggesting that they serve as a primary backbone structure supporting AM search processes. Moreover, results showed that the dlPFC was the only node in the network that served as an entry point for the external driving input; in the context of DCM, activity in entry point nodes is most prominently consistent with the main experimental manipulation, in this case, the onsets and durations of the AM search trials. The current results indicate that the dlPFC directly drives the activity of all other nodes in the network via positive connections during AM search. Even though the dlPFC has been to a certain extent associated with AM processes (Svoboda et al., 2006), that region is more often thought to play a cardinal role in higher-level executive functions related to cognitive control (Carlén, 2017), such as goal maintenance (Paxton et al., 2007). In recent years the dlPFC has become a remarkably common target in studies employing noninvasive brain stimulation techniques in a more general context of memory studies, where effects during both encoding and retrieval of episodic memories are examined, in both left and right hemispheres (often using the sites F3 and F4, respectively, under the 10-20 electroencephalogram system) (Chua et al., 2017; Gray et al., 2015; Habich et al., 2017; Manenti et al., 2018; Sandrini et al., 2013). The dlPFC has also been associated with processes underlying working memory (Curtis and D’Esposito, 2003), though more recent accounts attribute the activation of dlPFC in working memory studies to, again, mainly reflect cognitive control functions which are recruited during the execution of working memory tasks (Sreenivasan et al., 2014). This opens the possibility, as advanced by others (Nolde et al., 1998; Ranganath and Knight, 2002), that the involvement observed in the context of the AM search task was not entirely specific to the retrieval of episodic/autobiographical memories, but rather, majorly associated with functions pertaining to cognitive control or working memory processes that aim to fulfill the constraints and demands imposed by the experimental paradigm. One crucial aspect of such a role would be to coordinate episodic memory specific component processes taking place in the other nodes of the network. Though this does not necessarily preclude that the dlPFC has an essential function in the cued retrieval of AMs, the extent to which its contributions are specific to episodic memory retrieval processes still needs to be clarified by future research.

The DCM results also pointed out to the vmPFC as a major driver of activity within the network, with positive connections to all other nodes but the angular gyrus ROI. This result is in agreement with other studies that have highlighted the involvement of vmPFC in various stages of autobiographical memory processes (Barry et al., 2018; Barry and Maguire, 2019; Bonnici et al., 2012; Fuentemilla et al., 2014; McCormick et al., 2018; Nawa and Ando, 2019; Nieuwenhuis and Takashima, 2011) though in the context of episodic future imagining it has been reported larger effective connectivity from the hippocampus to the vmPFC (Campbell et al., 2017). A much theorized role of the vmPFC is related to the processing of memory schemas, i.e., knowledge representations about regularities found in typical contexts or experiences that are abstracted from multiple episodes, and which influence the acquisition and retrieval of memories (Gilboa and Marlatte, 2017; van Kesteren et al., 2012). Our result pointing out to a control exerted by the vmPFC on the hippocampus during AM retrieval is in line with the notion of temporal precedence of prefrontal regions over the hippocampus during retrieval of contextual representations (Place et al., 2016). Though the vmPFC and the dlPFC were linked by mutual positive connections, both connections were down-modulated during Hit trials, suggesting that both nodes partially disengage when a successful AM search occurs. Given the experimental context of this study, one straightforward interpretation of the current results is that, indeed, it is the vmPFC that largely coordinates processes specific to the retrieval of AMs, as opposed to a more general role played by the dlPFC. Still, what basic function would such a coordination actually involve? Here, the analysis focused in the period of time when participants were actively searching for a personal memory associated with a visually displayed cue. One possibility is that the vmPFC actively selects appropriate memories - in a broad sense - that fulfill a given criterion, in this case, the demand by the experimental task of memories having to be associated with the verbal cue. Studies with vmPFC patients have shown that a characteristic memory deficit displayed by vmPFC patients is *confabulation*, i.e., the retrieval of erroneous memories (Schneider and Koenigs, 2017). The current results are in agreement with such findings if considering that truthfulness is the most basic propriety of any *appropriate* memory.

The region in the posterior medial cortex, which here we deliberately labeled with a more general name (RSC/PCC/Prec), was positively influenced by the dlPFC, as all other nodes in the assessed network, but also by the vmPFC. In fact, the positive connections with the vmPFC went both ways, and were unaffected during Hit trials indicating that the vmPFC and the RSC/PCC/Prec were in lockstep during the search for AMs, regardless of the outcome. Interestingly, connections from the RSC/PCC/Prec to the dlPFC and dmPFC, which were negative on average across trials, were up-modulated during Hit trials (so much that the connection to the dlPFC achieved the greatest magnitude among all links), indicating that the posterior midline node likely plays a prominent role in the stages following successful AM search, most notably, elaboration.

The hippocampus has been historically viewed as a central structure supporting episodic memory, and it undeniably plays an essential role on various stages of episodic memory processes. However, there is a growing body of evidence showing that most memory processes should be conceptualized as being an interplay between the hippocampus and the prefrontal cortex (Eichenbaum, 2017). In agreement with that notion, results indicated that on average across trials of the AM search task, activity in the hippocampus was positively driven by the dlPFC and vmPFC. The hippocampus also played a major inhibitory role, negatively driving the activity of all other nodes in the network. Nevertheless, during Hit trials, the connections from the hippocampus to the vmPFC and the angular gyrus were up-modulated, signaling a more active involvement when AM search was successful. Previous studies have suggested functional and connectivity-wise differences along the anterior-posterior axis of the hippocampus (Blum et al., 2014; Chase et al., 2015; Fanselow and Dong, 2010; Zeidman and Maguire, 2016). The group-level peak voxel of the hippocampus ROI (MNI y = −28) used in the current study was located in a more posterior region of the hippocampus (Zeidman and Maguire, 2016). This could be a possible reason why we failed to observe an effect of the hippocampus in the dmPFC, as previously reported (McCormick et al., 2015). Studies have suggested the existence of anterior-posterior differences in the hippocampus in terms of the type of information that is retrieved (Nadel et al., 2013), and a memory advantage associated with the volume of the posterior hippocampus (Maguire et al., 2000; Poppenk and Moscovitch, 2011). Moreover, AM elaboration following successful AM search seems to recruit a more anterior region of the hippocampus (Nawa and Ando, 2019). All in all, these data suggest that there could be an anterior-posterior distinction between the two stages of AM retrieval. From a larger perspective, the current results suggest that the hippocampus is involved in the retrieval of AMs, even in the case of remote memories: the mean age of the memories was 2.8 years old (range [0.1 - 7.5]), indicating that for the large majority of participants in this study, the memories associated with the cues were likely to be more remote than recent.

Much like the hippocampus, results from the DCM analysis showed that the angular gyrus negatively drove the activity of the nodes with which it had effective connections (all but the dlPFC and vmPFC). However, in contrast to the hippocampus, during Hits, the negative connection with the RSC/PCC/Prec was further negatively enhanced, and most remarkably, two originally absent links with the dlPFC and vmPFC turned to become inhibitory connections. This pronounced involvement – albeit negative – with all other nodes during successful trials suggests that, like the hippocampus, the angular gyrus may play a more central role in subsequent stages of AM retrieval. Activity in the left angular gyrus scales with the recollection of fine-grained details from memory (Rugg and King, 2018), which has led some to advance the idea that the left angular gyrus has a fundamental role in the construction of perceptually rich imageries, irrespective of whether they are based on personal memories or hypothetical scenarios (Ramanan et al., 2017). A recent study employing noninvasive brain stimulation (Bonnici et al., 2018) showed a specific effect on the free recall of autobiographical memories (as opposed to the cued recall of AMs or the free or cued recall of word pairs) after inhibiting the activity in the left angular gyrus by means of a continuous theta burst stimulation: participants recalled fewer details of their AMs, plus fewer of the AMs were reported from a first-person perspective. These results causally implicate the left angular gyrus in the reconstruction of rich and detailed imageries of past experiences, which is the hallmark of AM recall or elaboration (Tulving, 1985; Wheeler et al., 1997). The inhibitory effect exerted by the angular gyrus specific during Hits in the three nodes that primarily drove the activity in the network during AM search (dlPFC, vmPFC and RSC/PCC/Prec) could be a reflection of the greater involvement of the angular gyrus during AM elaboration. An alternative hypothesis for the prominent inhibitory role of the angular gyrus in Hit trials would be along the lines of the “attention to memory” (AtoM) hypothesis regarding the involvement of lateral parietal regions in episodic memory retrieval processes (Cabeza et al., 2008; Ciaramelli et al., 2008). That model postulates a specific bottom-up attention control function to areas in the ventral parietal cortex (as opposed to the dorsal parietal cortex, which is thought to be involved with top-down attention processes) during the retrieval of memories, much in line with bottom-up attention control for sensory stimuli. One possible role of the ventral parietal cortex, including the angular gyrus, in the context of generative retrieval would be to drive attentional resources to internally generated relevant memory cues or retrieved memories in a bottom-up fashion. The inhibitory modulation driven by the angular gyrus in Hit trials could thus be interpreted as signaling the termination of the generative retrieval process due to the successful completion of memory search. Other models have been proposed to explain the involvement of the lateral parietal cortex in episodic memory processes (Sestieri et al., 2017), clearly further work is necessary to clarify this question.

Corroborating the DCM findings, functional connectivity analyses based on resting-state data (see Supporting Information) collected before the AM search task from the same participants confirmed that the vmPFC, hippocampus, angular gyrus and RSC/PCC/Prec form of a tightly knit clique, even when a task was not externally imposed. On the other hand, the dlPFC and the dmPFC were only connected to the other nodes via the angular gyrus.

This study has a few caveats that must be kept in mind when interpreting these results. First and foremost, DCM results were obtained using an approach that relied on the automatic pruning of fully-connected models, which though principled, explored a model space that is possibly much larger than what is typically assessed in purely hypothesis-driven DCM studies. This could possibly complicate the interpretation of the results; just like with any other experimental result, they still have to withstand the test of replicability. Another limitation of this study is that the current 6-node network is obviously not an exhaustive representation of all brain regions that have been associated with episodic memory retrieval processes; therefore, the possibility that left out regions may have indirectly influenced the observed dynamics, or that their inclusion may qualitatively and quantitatively alter the interactions described here cannot be categorically ruled out. Also, the DCM results presented here were based on an indirect, low-temporal resolution measure of brain activity (BOLD), with all its merits and limitations. This drawback could certainly be better resolved in the future by means a combination of different neuroimaging modalities, such as magnetoencephalography (Barry et al., 2019b; Garrido et al., 2015).

In summary, though there is still much work to be done to more comprehensively and accurately characterize the functions and computations performed by the various brain regions recruited in the performance of human episodic memory capacities, first and foremost, the current results highlight the interaction of several brain regions during the cued search for AMs. More specifically, these results suggest that midline cortical regions together with the dlPFC are the initiators of AM search process, setting up the basic structure on which the angular gyrus and the hippocampus can act upon when the outcome of the search is successful. Most importantly, they indicate that the interplay among these regions is what enables navigating through our memories and that indeed, targeting such interactions might be the shortest path to ameliorate memory disorders that affect many.

## Acknowledgements

We are indebted to the imaging staff at the Center for Neural and Information Neural Networks for their invaluable and often ingenious help running the imaging experiments. This research was supported by the Japan Society for the Promotion of Science under a Grant-in-Aid for Scientific Research (JSPS KAKENHI grant number JP17K00220 to NEN). The funding source was not involved in any stage of the research described in this article.

