## Supporting Information for "Effective connectivity during autobiographical memory search"

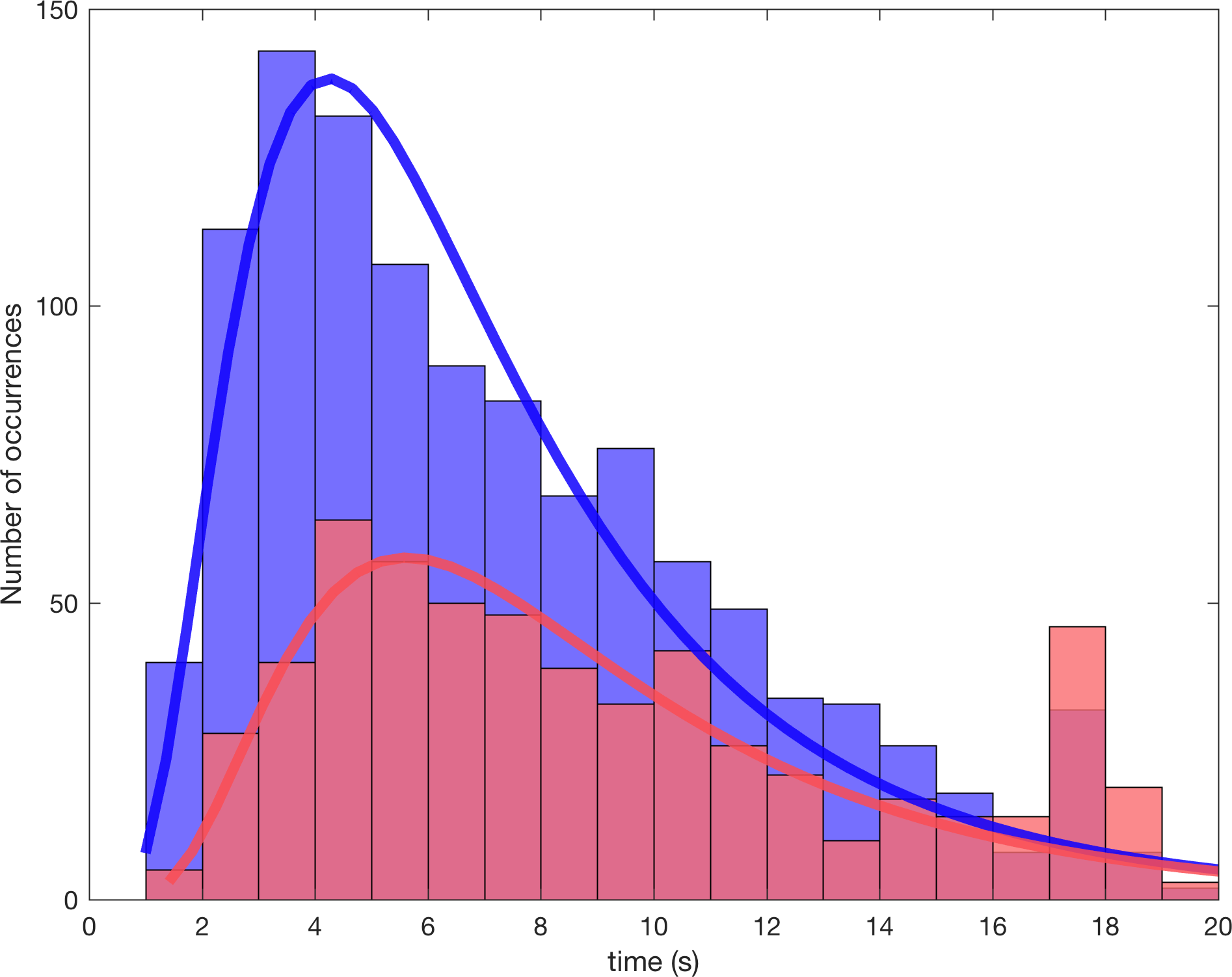

**FIG S1:** Histograms showing the reaction times for the Hit trials (blue) and the Miss trials (red), for the 24 participants included in the analyses. The continuous lines show the best-fit lognormal distributions based on the same data. Seventeen seconds after the onset of the verbal cue (t = 0), the color of the cue changed to purple to signal participants that there were only 3 seconds remaining until the end of the trial. In the Miss trials, a large portion of the button presses were generated right after the cue displayed on the screen that served to guide the AM search changed colors to signal that the end of the trial was approaching (third red bin from the right, corresponding to the period of the time 17 to 18 seconds after cue onset).

**Section S1. Preprocessing of resting-state data**

The preprocessing pipeline for the resting-state data started with the realigned and unwarped data, and continued as follows: detection of outlier scans based on the Artifact Detection Tools (ART, https://www.nitrc.org/projects/artifact_detect, RRID: SCR_005994), segmentation of mean functional image and anatomical image into grey matter, white matter and cerebrospinal fluid (CSF) masks, and normalization to MNI standard brain, and finally, spatially smoothing of functional images using a 6 mm FWHM Gaussian kernel. Following that, the default denoising protocol implemented in the CONN toolbox was applied to the data, the anatomical CompCor (Behzadi et al., 2007), which addresses global-level artifacts embedded in voxelwise timeseries by computing representative white matter and CSF components and treating them as nuisance parameters, together with movement parameters and outlier scans. The residuals derived from this procedure were then band-pass filtered to 0.008-0.09 Hz, and the filtered data were used in the resting-state functional connectivity analysis. Both DCM and resting-state functional connectivity analyses were performed using data normalized to the Montreal Neurological Institute (MNI) template space data.

**Section S2. Examining functional connectivity in the AM retrieval network during resting-state**

We examined the underlying functional connectivity between the ROIs by performing a ROI-to-ROI analysis and a ROI-to-voxel analysis based on the data from the resting-state session. The ROIs were 5 mm-spheres centered at the same group-level peak voxels used in the DCM analysis. ROI timeseries were the averaged voxelwise timeseries of all voxels within the spheres. A ROI-to-ROI analysis assesses how the activity in specific regions of interest temporally covariate with one another. Bivariate correlations (Pearson’s r) were computed between ROI timeseries, for all participants, and the results for each pair of ROIs were entered in a one-sample t-test after being Fisher Z-transformed. The results from the t-tests were examined for statistical significance at a level of p < 0.05 using false discovery rate (FDR) correction (Chumbley et al., 2009) at the analysis level, i.e., taking into account the total number of ROIs used in the analysis. A ROI-to-voxel analysis, on the other hand, assesses more broadly how the activity in a given ROI covariates with regard to all other voxels in the brain. The bivariate correlation between the ROI timeseries and all other voxels in the brain were computed, for all participants. Voxelwise r values for each participant were entered in a one-sample t-test after being Fisher Z-transformed. Results were examined for statistical significance at a voxelwise threshold level of p < 0.001 uncorrected, and a cluster-level threshold of p < 0.05, using FDR correction.

**Section S3. Resting-state data functional connectivity analysis**

Resting-state data were available for 21 participants. The 3 participants from whom we could not collect resting-state data were included in the DCM analysis cohort. Note that the resting-state data was collected prior to the performance of the AM search task. The group-level peak voxels used in the DCM analysis (Table 2) were used to determine the locations of the ROIs (5 mm spheres centered at each peak voxel). Note that differently from the DCM analysis, no individual level adjustments were performed; timeseries were collected from the same MNI coordinates for all participants. Results for the ROI-to-ROI analysis are summarized in Figure S2. The vmPFC, hippocampus, angular gyrus and RSC/PCC/Prec formed a full-connected clique. In contrast, the dmPFC was only connected to the dlPFC, which on its turn was connected to the angular gyrus. These results were to a great extent replicated by the ROI-to-voxel analysis (Figure S3). Largely overlapping patches of highly connected voxels were observed when using the vmPFC, hippocampus, angular gyrus and RSC/PCC/Prec as seeds. Moreover, when using the dlPFC as a seed, patches overlapping with the angular gyrus ROI and the dmPFC ROI were observed, though connectivity with the other ROIs in the midline (vmPFC, hippocampus, and RSC/PCC/Prec) was inexistent. When using the dmPFC ROI as a seed, connectivity with an area overlapping with the dlPFC ROI was observed. Curiously, though no overlap with the angular gyrus ROI was observed – replicating the ROI-to-ROI findings – there was a patch of connectivity with a more lateral and anterior region of the parietal cortex than the angular gyrus ROI.

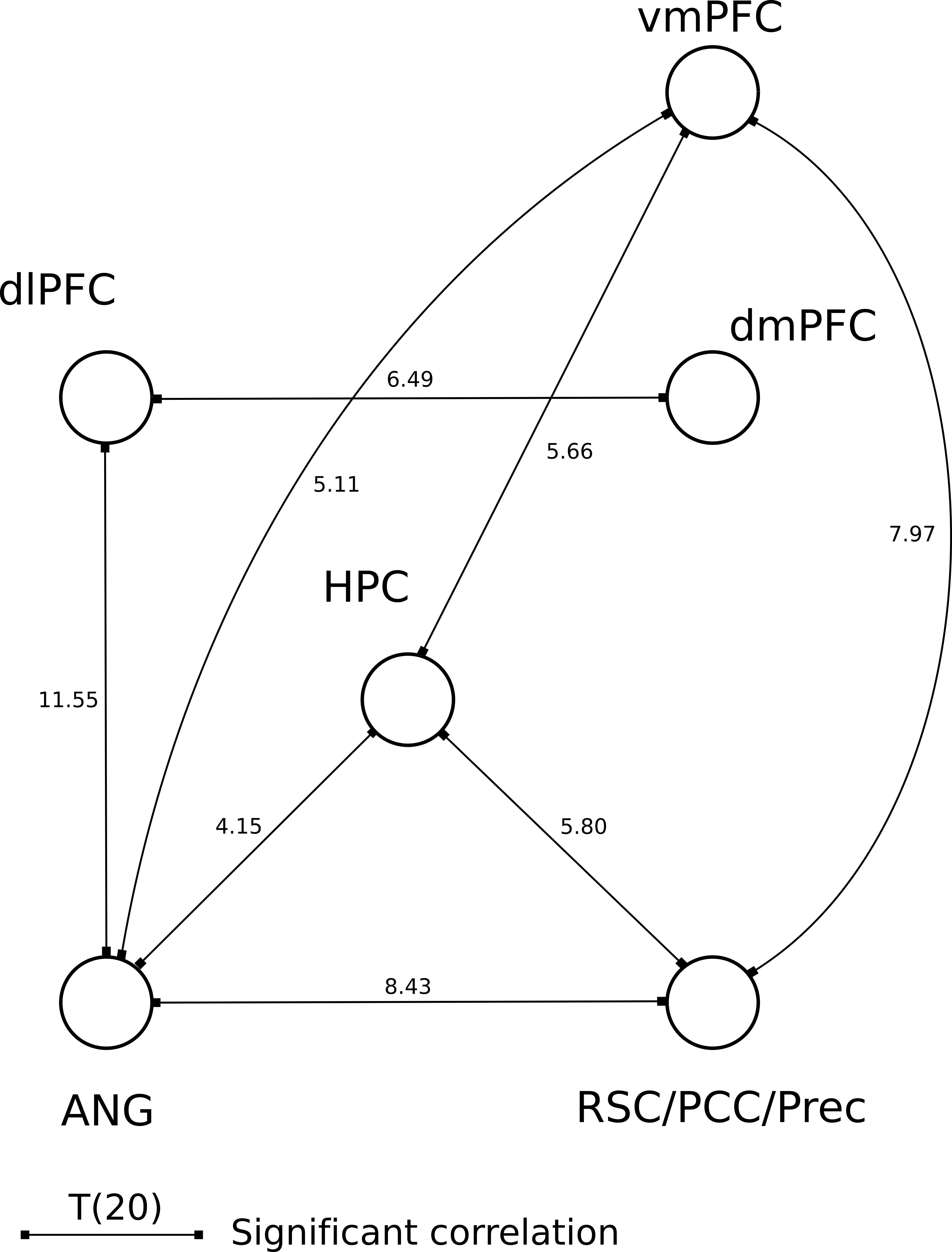

**Figure S2:** *Results of the ROI-to-ROI analysis. Only the connections that survived a p < 0.05 FDR correction at the analysis level are displayed (with the corresponding t-test statistic).*

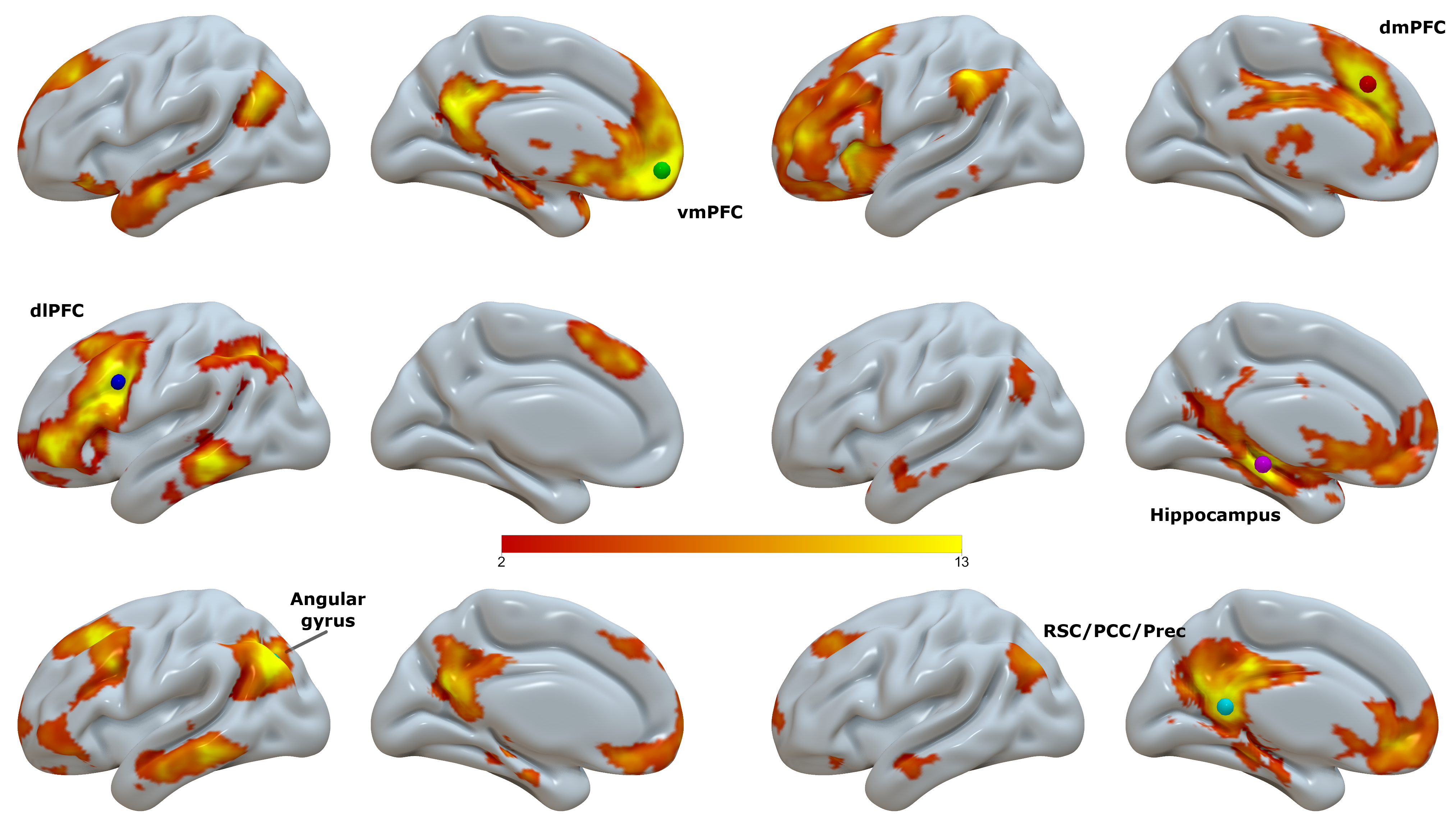

**Figure S3:** *Panels showing the group-level results of the ROI-to-voxel functional connectivity analysis, for each one of the 6 ROIs (locations displayed as colored spheres). T-values are overlaid on a semi-inflated smoothed version of the ICBM152 brain using Surf Ice. Results are shown at a voxelwise threshold level of p < 0.001 uncorrected, and a cluster-level threshold of p < 0.05, using FDR correction. Only the results for the positive contrast are shown (i.e., voxels that were negatively correlated with the respective ROIs are not displayed).*
